# Scrutiny of human lung infection by SARS-CoV-2 and associated human immune responses in humanized mice

**DOI:** 10.1101/2021.11.05.466755

**Authors:** Renren Sun, Zongzheng Zhao, Cong Fu, Yixin Wang, Zhendong Guo, Chunmao Zhang, Lina Liu, Cheng Zhang, Chang Shu, Jin He, Shucheng Hua, Yuwei Gao, Zheng Hu

## Abstract

There is an urgent need for animal models of COVID-19 to study immunopathogenesis and test therapeutic intervenes. In this study we showed that NSG mice engrafted with human lung (HL) tissue (NSG-L mice) could be infected efficiently by SARS-CoV-2, and that live virus capable of infecting Vero cells was found in the HL grafts and multiple organs from infected NSG-L mice. RNA-seq examination identified a series of differentially expressed genes, which are enriched in viral defense responses, chemotaxis, interferon stimulation, and pulmonary fibrosis between HL grafts from infected and control NSG-L mice. Furthermore, when infecting humanized mice with human immune system (HIS) and autologous HL grafts (HISL mice), the mice had bodyweight loss and hemorrhage and immune cell infiltration in HL grafts, which were not observed in immunodeficient NSG-L mice, indicating the development of anti-viral immune responses in these mice. In support of this possibility, the infected HISL mice showed bodyweight recovery and lack of detectable live virus at the later time. These results demonstrate that NSG-L and HISL mice are susceptible to SARS-CoV-2 infection, offering a useful *in vivo* model for studying SARS-CoV-2 infection and the associated immune response and immunopathology, and testing anti-SARS-CoV-2 therapies.

## Introduction

The lack of animal models that can be easily used to model SARS-CoV-2 infection and pathogenesis in biosafety level 3/4 facilities is an important drawback factor impeding mechanistic understanding of coronavirus disease 2019 (COVID-19) pathogenesis and test new anti-viral therapies(1). Although mice are the most popular and easily handleable animal model, conventional murine models are not suitable for COVID-19 studies because most SARS-CoV-2 strains cannot use murine angiotensin-converting enzyme 2 (ACE2) to invade mouse cells(2). Although human ACE2 transgenic mice have been shown susceptible to SARS-CoV-2 infection(3, 4), this model overlooks other proteins involved, such as transmembrane serine protease 2 (TMPRSS2) and CD147(5, 6). In addition, profound evolutionary divergences between mice and humans also compromise their value to faithfully replicate viral infection process and the associated disorders(7). Non-human primates are genetically close to human and can be infected by SARS-CoV-2(8), however, their wide usage is intensively restrained due to their extreme high cost and the need for complicated operating procedures especially in biosafety level 3/4 facilities. Thus, there is an urgent need for novel animal models that are easy to handle and faithfully mimic human SARS-CoV-2 infection and the associated immune responses.

Immune surveillance plays crucial roles in the control of viral infection and the development of inflammatory syndromes, including pneumonia and cytokine storm in COVID-19 patients(9). Previous studies of ours and other groups have shown that human immune system (HIS) mice made by co-transplantation of human fetal thymic tissue (under renal capsule) and CD34^+^ fetal liver cells (FLCs, i.v.)(10, 11) could mount antigen specific human T and antibody responses following immunization, viral infections or transplantation(12–14). Recently, we generated a HIS mouse model with autologous human lung (HL) tissues (referred to as HISL) by transplantation of both human fetal lung (subcutaneous) and thymic tissues and CD34^+^ FLCs, and validated its efficacy to study human lung resident immunity and anti-viral response for H1N1 viral infection(15). Herein, we further explored its application for SARS-CoV-2 infection investigation. We found that these HISL mice could be infected by SARS-CoV-2 virus and develop anti-viral immune responses, offering a convenient and useful *in vivo* model for understanding COVID-19 pathogenesis and testing anti-SARS-CoV-2 interventions.

## Results and Discussion

### Entry and replication of SARS-CoV-2 in HL xenografts in mice

We first determined whether human lung tissue grafted in NOD/SCID IL2rg^-/-^ (NSG) mice can be infected by SARS-CoV-2. Briefly, NSG mice were implanted subcutaneously with HL tissue (NSG-L mice), and subjected to infection via intra-HL injection of SARS-CoV-2 or PBS (as controls) 6-8 weeks later. Immunohistochemistry (IHC) examination confirmed human ACE2 protein expression throughout the HL xenografts (Figure 1A). Real-time qPCR (RT-qPCR) analysis revealed that SARS-CoV-2 RNA was detected in HL and various mouse tissues including heart, liver, spleen, lung, kidney, brain and intestine at days 1 and 3 following inoculation of 10^6^ TCID_50_ SARS-CoV-2, but not in NSG mice receiving subcutaneous injection of a similar number of virus (Figure 1B), demonstrating SARS-CoV-2 infection and expansion in HL xenografts. To confirm this observation, another cohort of NSG-L mice was intra-HL inoculated with SARS-CoV-2 or mock. Again, viral copies were detected in HL at days 3, 5, 8, 14 and 21 after infection (Figure 1C). Importantly, 13 out of 14 HL homogenate samples (except one at day 21) harvested from SARS-CoV-2-infected NSG-L mice at days 3, 5, 8, 14, and 21 after infection retained the capability to re-infect Vero E6 cells *in vitro*, demonstrating the existence of live SARS-CoV-2 in HL xenografts (Figure 1D). However, viral inoculation did not lead to significant bodyweight loss (Figure 1E) or detectable pathological changes in HL grafts (Figure 1F) or mouse tissues (Supporting Information Figure 1), likely due to the immunodeficiency of NSG-L mice. This is in line with previous reports that the complications in COVID-19 patients are largely attributable to immune responses driven by SARS-CoV-2 infection(16).

**Figure 1.**
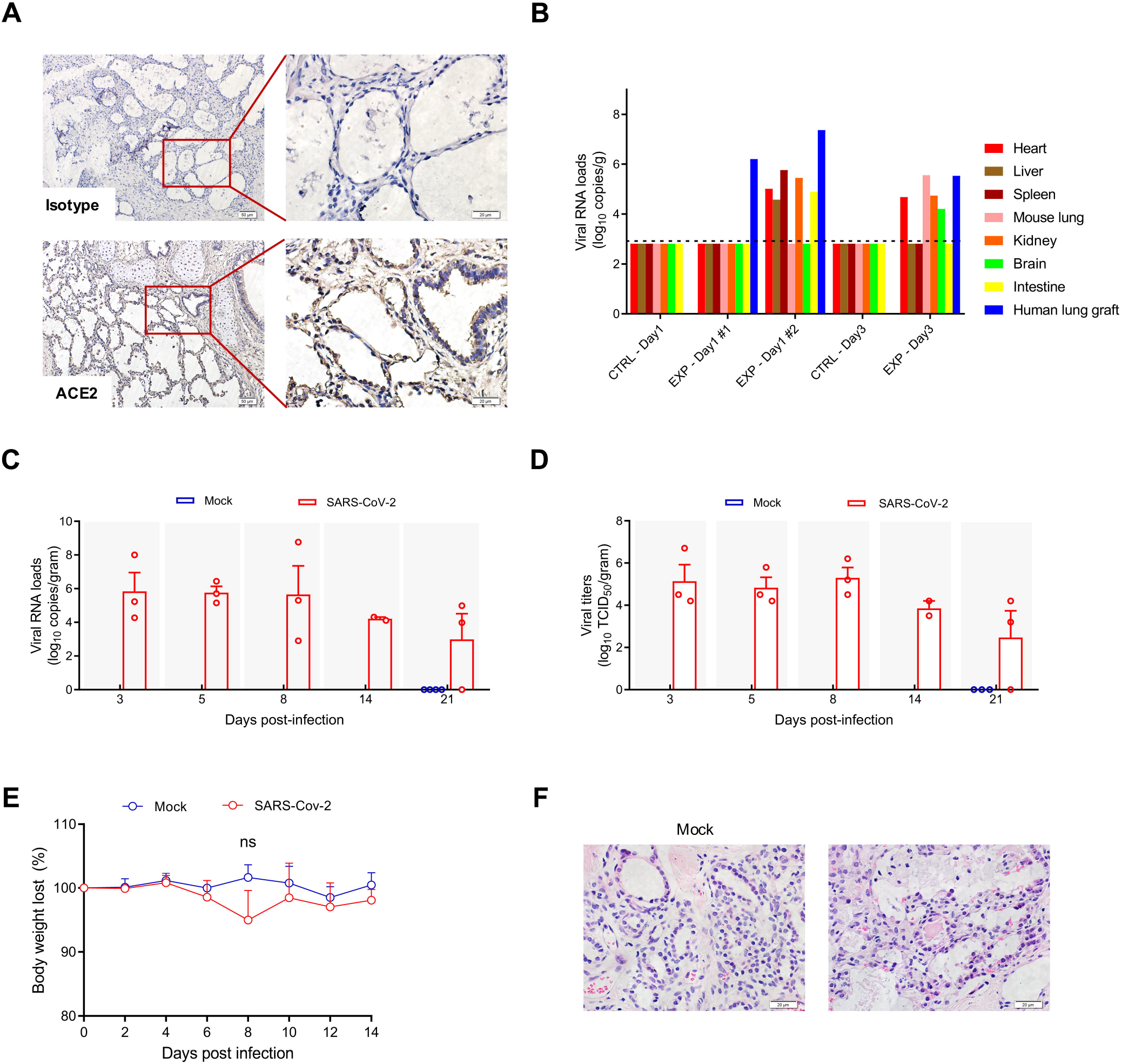
SARS-CoV-2 infection of HL xenografts in NSG mice. NSG-L mice were subjected to SARS-CoV-2 infection (10^6^ TCID_50_ given by intra-HL injection) 6-8 weeks after HL transplantation. Data from two independent experiments are presented, in which NSG mice receiving subcutaneous injection of SARS-CoV-2 (**B**) or NSG-L mice receiving intra-HL injection of PBS (**C-F**) were used as controls, respectively. **(A)** Representative IHC images of HL xenografts (up, isotype; down, human ACE2). **(B)** Viral copies in the indicated tissues examined by RT-qPCR 1 (n=2 and 1 for HL-infected and control groups, respectively) or 3 (n=1 per group) days after infection. **(C-F)** Viral RNA copies examined by RT-qPCR (**C**) and titers measured by Vero E6 cells (**D**) in HL xenografts, and bodyweight changes (**E**) at the indicated time points (n=3 per group); (**F**) Representative H&E images of HL xenografts at day 5 after intra-HL injection of PBS (left) or SARS-CoV-2 (right).

### RNA-seq analysis for HL xenografts after SARS-CoV-2 infection

To further understand the insight into SARS-CoV-2 infection induced changes in HL cells, we analyzed the transcriptome by RNA-seq of HL samples from uninfected NSG-L mice or infected NSG-L mice 5 and 21 days after SARS-CoV-2 inoculation. For day-5 HL grafts, a total of 4612 human genes were differentially expressed between two samples, of which 2129 genes were significantly upregulated and 2483 genes were downregulated (|*log2(FoldChange)*| > *1 & padj< 0.05*) in SARS-CoV-2-infected compared to uninfected HL grafts (Figure 2A left). For day-21 HL grafts, a total of 6488 human genes were differentially expressed between the two samples, of which 3311 and 3177 genes were significantly upregulated and downregulated (|*log2(FoldChange)*| > *1 & padj< 0.05*), respectively in SARS-CoV-2-infected compared to uninfected HL grafts (Figure 2A right). Many of these differentially expressed genes are associated with viral defense responses, including NLRC5, MICB, APOBEC3D, APOBEC3G, IFI6, ISG15, and IFITM3, which were markedly upregulated in infected HL grafts at days 5 and 21 compared to HL grafts from uninfected NSG-L mice (Figure 2B). Notability, HL grafts from the infected NSG-L mice showed a great upregulation of viral infection-associated chemokines (e.g., CCL11 and CXCL6 at day 5, and CCL19, CXCL19, CXCL13, CCL18 at day 21; Figure 2C) and interferon-stimulated genes (ISGs; e.g., IFIT1 and STAT1 at day 5, and ISG15, IFITM1 and IFI27 at day 21; Figure 2D) compared to those from non-infected NSG-L mice. Furthermore, SARS-CoV-2 infection induced intensive upregulation of genes involved in pulmonary fibrosis at day 5 (e.g. CSF1, FASLG, NLRP1) and day 21 (e.g. IGF2, IL18, MMP9) in infected HL grafts (Figure 2E), providing a mechanistic explanation for the reported development of lung fibrosis, a long-term and presumably irreversible complication of COVID-19 patients(17). Together, these findings support the use of NSG-L mice as a convenient and useful *in vivo* platform for modeling human lung infection with SARS-CoV-2 and testing anti-viral therapies.

**Figure 2.**
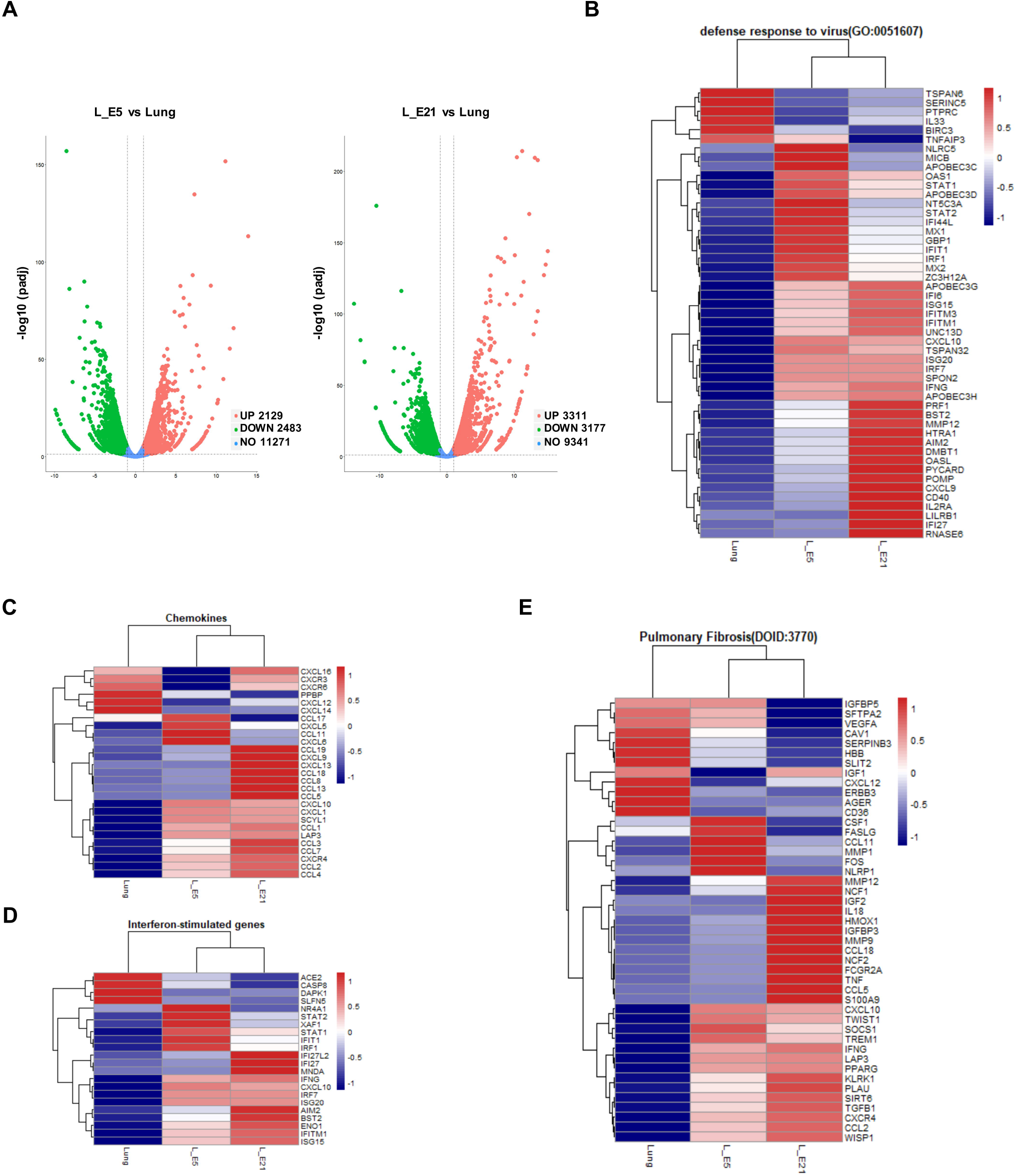
RNA-seq analysis for HL grafts from NSG-L mice after SARS-CoV-2 infection. HL grafts injected with 10^6^ TCID_50_ SARS-CoV-2 virus were harvested from NSG-L mice for RNA-seq examination at day 5 and 21 after infection; Naïve HL graft without infection was used as control. **(A)** Volcano plot showing DEGs of HL grafts for SARS-CoV2-infected at day 5 (left) or day 21 (right) and non-infection control from NSG-L mice. The data for all genes are plotted as log2 fold change versus the −log10 of the adjusted p-value. **(B-E)** Heatmaps of RNA-seq expression Z-scores computed for selected genes that are differentially expressed (*p adj* < 0.05, |log 2 (foldchange)| > 1) between 3 pairwise comparisons (L_E5 vs L_E21, Lung vs L_E5, Lung vs L_E21). (**B**) DEGs under GO term “defense response to virus” (BP GO: 0051607); (**C**) Chemokines; (**D**) Interferon-stimulated genes; **(E)** DEGs under DO term “Pulmonary Fibrosis” (DOID:3770).

### Development of antiviral immunity in HISL mice following SARS-CoV-2 infection

We next examined SARS-CoV-2 infection in HISL mice with both HL and human immunity. Briefly, HISL mice were constructed by transplantation of HL and thymic tissues and CD34^+^ cells, and subjected to SARS-CoV-2 infection after human immune reconstitution was confirmed by measuring human immune cells in peripheral blood. FACS analysis of PBMCs 14 weeks after humanization detected high levels of human CD45^+^ lymphohematopoietic cells composed of CD3^+^ T cells (including both CD4 and CD8 T cells), CD20^+^ B cells and CD33^+^ myeloid cells in HISL mice (Supporting Information Figure 2A). Furthermore, human CD45^+^ immune cells including CD4^+^ T cells, CD20^+^ B cells and CD11c^+^ dendritic cells were also detected in spleen and HL grafts from these mice (Supporting Information Figure 2B). These HISL mice were infected 3 weeks later with SARS-CoV-2 or mock as controls. Unlike NSG-L mice (Figure 1), HISL mice showed a significant bodyweight loss after intra-HL SARS-CoV-2 infection, which returned to a level comparable to the control HISL mice by 21 days (Figure 3A). No mortality was observed in HISL mice infected with SARS-CoV-2. Furthermore, HL grafts from HISL mice that received SARS-CoV-2 (but not mock) had hemorrhage at day 5, which was associated with increased infiltration by human CD45^+^cells, including CD4^+^ T and CD20^+^ B cells (Figure 3B). Although the levels of human T and B cell infiltration in HL grafts became comparable between the two groups at day 21, there were still significantly more human CD33^+^ myeloid cells (Supporting Information Figure 3A), including CD11c^+^ cells (Supporting Information Figure 3B), in HL grafts in SARS-CoV-2-infected than mock infected mice, which is in line with previous reports for patients(18, 19). No significant difference was detected for any human immune cell populations in WBCs, including CD3^+^ T, CD20^+^ B, CD33^+^ myeloid, and CD56^+^ NK cells between SARS-CoV-2- and mock-infected mice at day 21 (Supporting Information Figure 3C). Viral copies were detected by RT-qPCR in all HL samples (except one at day 21) from SARS-CoV-2-infected HISL mice, but not those from the control HISL mice (Figure 3C). However, in contrast to HL homogenates from infected NSG-L mice (Figure 1), HL homogenate samples from most infected HISL mice harvested at day 5 (5 out 6) and day 21 (5 out 6) failed to infect Vero E6 cells *in vitro* (Figure 3D), indicating an efficient elimination of live SARS-CoV-2 by human immune system in these mice. Together, these results suggest that the clinical symptoms were caused mainly by inflammation/immune responses, but not viral invasion directly, supporting the use of HISL mice to study COVID-19 pathogenesis and anti-SARS-CoV-2 immunotherapy.

**Figure 3.**
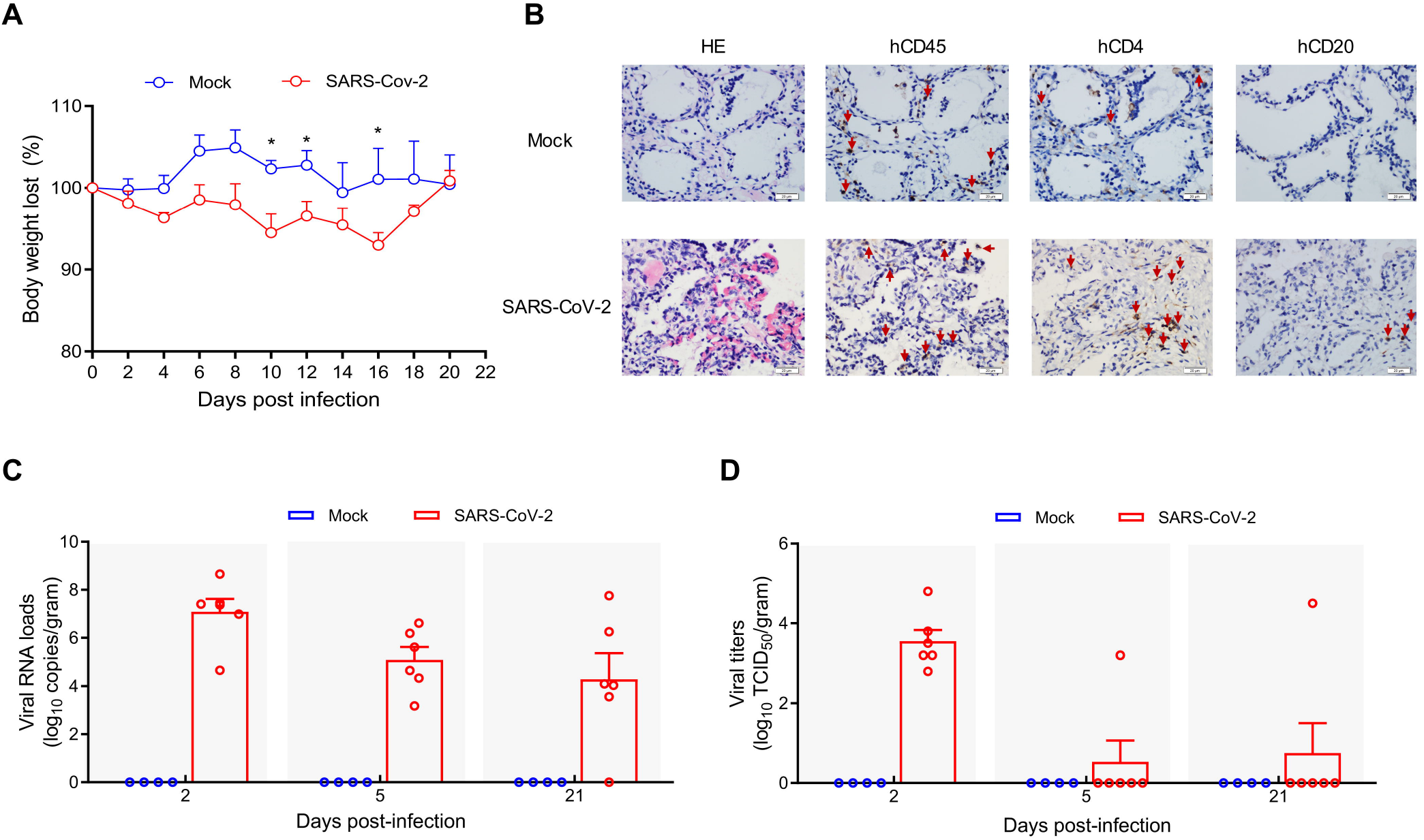
HISL mice are susceptible for SARS-CoV-2. HISL mice were infected at week 17 post-humanization with mock or 10^6^ TCID_50_ SARS-CoV-2 into HL. (**A**) Bodyweight changes of HISL mice infected with SARS-CoV-2 (n=6) or mock (n=4). (**B**) Representative H&E and IHC images of HL sections of HISL mice infected with SARS-CoV-2 (n=6) or mock (n=4) at day 5. (**C-D**) Mice were euthanized at days 2, 5, and 21 post-infection, and SARS-CoV-2 viral copies (**C**) and viral titers (**D**) in HL were examined (each symbol represents an individual animal). \**P* < 0.05.

### Concluding Remarks

In this study, we established a humanized mouse model carrying human lung tissue and autologous immune system, which can be easily constructed and comfortably used in ABSL-3 laboratories. We found that these humanized mice are susceptible to infection by SARS-CoV-2 and develop disease-associated lung injury. Although not capable of modeling upper respiratory or systemic infection/manifestations, as only the HL xenograft is susceptible to SARS-CoV-2 in these mice, this model is still useful as the only currently available model that permits *in vivo* assessment of human immune responses against SARS-CoV-2 infection. Therefore, the HISL humanized mouse model offers a convenient and powerful *in vivo* model for evaluating the immunopathology of COVID-19 and the efficacy of anti-viral immunotherapies, such as vaccination.

## Material and methods

### Mice and human tissues

NOD-Prkd^cem26Cd52^Il2rg^em26Cd22^/Nju (referred to as NSG) mice were purchased from Nanjing Biomedical Research Institute of Nanjing University. All mice were housed in a specific pathogen-free (SPF) microisolator environment and used between 6-8 weeks of age. Discard human fetal samples of gestational age of 17-22 weeks were obtained with informed consent at the First Hospital of Jilin University. Human fetal thymic and lung tissues were cut into small pieces with a diameter of around 1 mm or 5 mm, respectively, and CD34^+^ cells were purified from fetal liver cells (FLCs) by magnetic-activated cell sorting (MACS; with a purity of >90% confirmed by FACS). The human tissues and cells were cryopreserved in liquid nitrogen until use.

### NSG-L mouse and HISL humanized mouse model construction

NSG-L mice were made by subcutaneous implantation of human fetal lung tissue (approximately 5 mm in diameter). HISL humanized mice were made by implantation of human fetal thymic tissue (around 1 mm in diameter; under the renal capsule) and lung tissue (5 mm in diameter; subcutaneously) in NSG mice that pre-conditioned with 1.75 Gy total body irradiation (TBI; Rad Source RS2000-225, USA), followed by intravenous injection of 1-2×10^5^ human CD34^+^ FLCs (given 6-8 hours after TBI).

### Flow cytometric analysis

Mouse PBMCs were prepared using density gradient centrifugation with Histopaque 1077 (Sigma-Aldrich, St. Louis, MO). Human immune cell reconstitution in humanized mice were determined by flow cytometry using following fluorochrome-conjugated antibodies: anti-human CD45, CD3, CD4, CD8, CD20, CD33, CD56; anti-mouse CD45 and Ter119 (BD Biosciences, USA). Examination was performed on a Cytek Aurora (Cytek Biosciences, USA) and data were analyzed using the FlowJo software version 10.6.2. Dead cells were excluded from the analysis by gating out propidium iodide retaining cells.

### SARS-CoV-2 virus preparation and titer determination

SARS-CoV-2 (BetaCoV/Beijing/IME-BJ05-2020) was proliferated in Vero E6 cells, which are maintained in Dulbecco’s modified Eagle’s medium (DMEM; Invitrogen, Carlsbad, CA, USA) with supplemented 2% fetal bovine serum (FBS; Gibco, Auckland, New Zealand). Viral titers were determined using a standard 50% tissue culture infection dose (TCID_50_) assay.

### Mouse experiments

NSG-L or HISL mice were anesthetized with isoflurane and injected with 100 μl of 10^6^ TCID_50_ of SARS-CoV-2 or PBS into the human lung (HL) graft. The mice were followed for bodyweight changes, and euthanized for measuring viral particles and histological changes in HL grafts and tissues at indicated time points.

### RNA extraction and RT-qPCR

Total RNA was extracted from tissues homogenates using the RNeasy Mini Kit (Qiagen, Hilden, Germany), and viral RNA loads were determined by a SARS-CoV-2 RNA detection kit (Shenzhen Puruikang Biotech, China). The viral RNA loads for the target SARS-CoV-2 N gene was normalized to the standard curve obtained by using a plasmid containing the full-length cDNA of the SARS-CoV-2 N gene. The reactions were performed with CFX96 system (BIO-RAD, USA) according to the following protocol: 50 °C for 20 min for reverse transcription, followed by 95 °C for 3 min and then 45 cycles of 95 °C for 5 s, 56 °C for 45 s. Results were presented as log10 numbers of genome equivalent copies per gram of sample.

### Histology

Tissues were embedded in paraffin, and sectioned (2.5 μm) for H&E and immunohistochemistry (IHC) examination. For IHC, tissue sections were firstly stained with monoclonal anti-human CD45 (DAKO, 2B11+PD7/26), CD20 (DAKO, L26), CD4 (ABclonal; ARC0328), or CD11c (Abcam; EP1347Y), ACE2 (Abcam; EPR4435(2)) antibodies, and the immunoreactivity was detected with UltraSensitiveTM Streptavidin-Peroxidase Kit (KIT-9710, Mai Xin, China) according to the manufacturer’s protocol.

### RNA-seq

Total RNA was polyA-selected, fragmented and the samples were examined on an Illumina Novaseq6000 machine, yielding an average of 22 million uniquely aligned paired-end 150-mer reads per sample. Reads were aligned with STAR RNA-seq aligner (Version 2.7.3a) using the UCSC/hg38 genome assembly and transcript annotation(20). Expression levels were calculated as Fragments per Kilobase of transcript per Million reads (FPKM) using Cufflinks software(21). Differential expression analysis was performed using Cuffdiff (version 2.2.1.Linux_x86_64)(21). In each analysis, genes with thresholds of |log 2 (foldchange)| > 1, *p adj* < 0.05 and a mean FPKM value of more than 5 were tested. “Cluster Profiler” package (Release 3.16.1)(22) and “DOSE” package (Release 3.14.0)(23) in R software (version 4.0.2) was used for functional enrichment analysis, and GO biological processes terms at the significant level (*q-value* < 0.05) were employed.

### Statistical analysis

Data were analyzed using GraphPad Prism 8 software (San Diego, CA, USA) and presented as mean values ± SEM. The results were compared statistically using unpaired two-tailed Student’s t test and two-way analysis of variance (ANOVA), and a p value of ≤ 0.05 was considered significant.

## Abbreviations

HL: human lung
NSG-L: NSG mice engrafted with human lung tissue
HIS: humanized mice with human immune system
HISL: humanized mice with human immune system and autologous human lung
SARS-CoV-2: severe acute respiratory syndrome coronavirus 2
COVID-19: coronavirus disease 2019
ACE2: angiotensin-converting enzyme 2
IHC: Immunohistochemistry
RT-qPCR: Real-time qPCR
RNA-seq: RNA sequencing
DEGs: differentially expressed genes
DO: disease ontology
GO: gene ontology

## Data availability statement

RNA-seq data have been deposited into the National Center for Biotechnology Information (NCBI) Gene Expression Omnibus (GEO) database under accession numbers GSE175900.

## Ethics approval statement for human and animal studies

Protocols involved in the use of human samples and animals were reviewed and approved by the Institutional Review Board and Institutional Animal Care and Use Committee of the First Hospital of Jilin University, and all experiments with SARS-CoV-2 (COVID-19) were performed in biosecurity level 3 laboratory according to the protocols approved by the Changchun Veterinary Research Institute, Chinese Academy of Agricultural Sciences.

## Author contributions

Z.H., Y-W. G., S-C.H. conceived the study; R-R.S., Z-Z.Z., Y-X.W., C.S., J.H. performed the experiments; C.F. contributed to RNA-seq data analysis; Z.H., Y-W.G., S-C.H. supervised the work; Z.H., R-R.S., Z-Z.Z., C.F. analyzed the data and wrote the manuscript; all authors edited and approved the manuscript.

## Acknowledgements

This work was supported by grants from the Strategic Priority Research Program of the Chinese Academy of Sciences (XDA16030303), Chinese MOST (2017YFA0104402) and NSFC (91742107, 81870091, 91642208, 82001697 and 81941008), Science development project of Jilin province (20190201295JC and 20200703012ZP), and special grants for COVID-19 research of Jilin province (20200901006SF).

