## Supplemental Figures for "Scrutiny of human lung infection by SARS-CoV-2 and associated human immune responses in humanized mice"

### Supporting Information

Supporting Information Figures: 4

A B C D E F

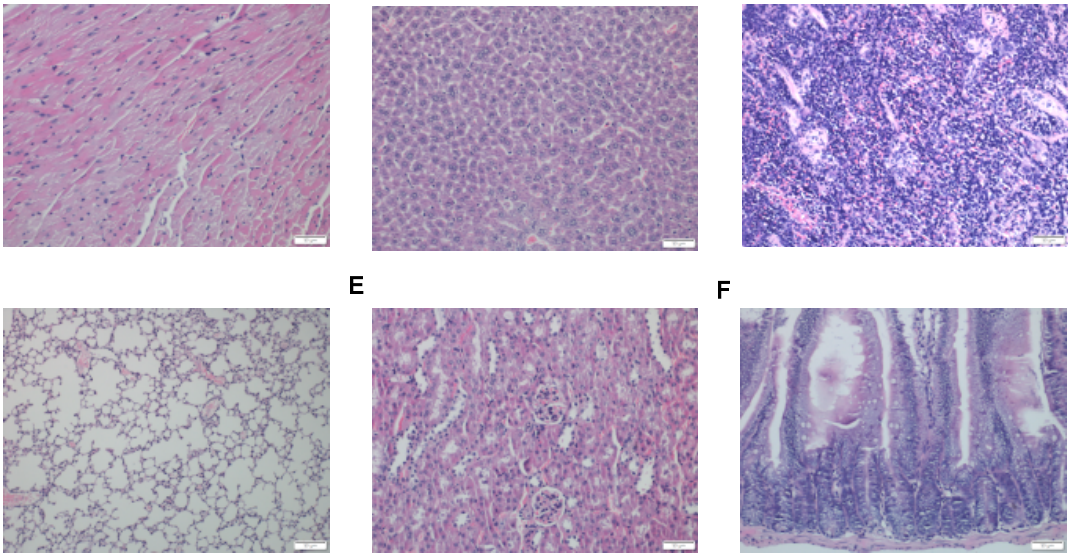

**Supporting Information Figure 1.** Representative H&E-stained sections of heart **(A)**, liver **(B)**, spleen **(C)**, lung **(D)**, kidney **(E)** and small intestine **(F)** harvested from NSG-L mice at day 5 post-infection with SARS-CoV-2.

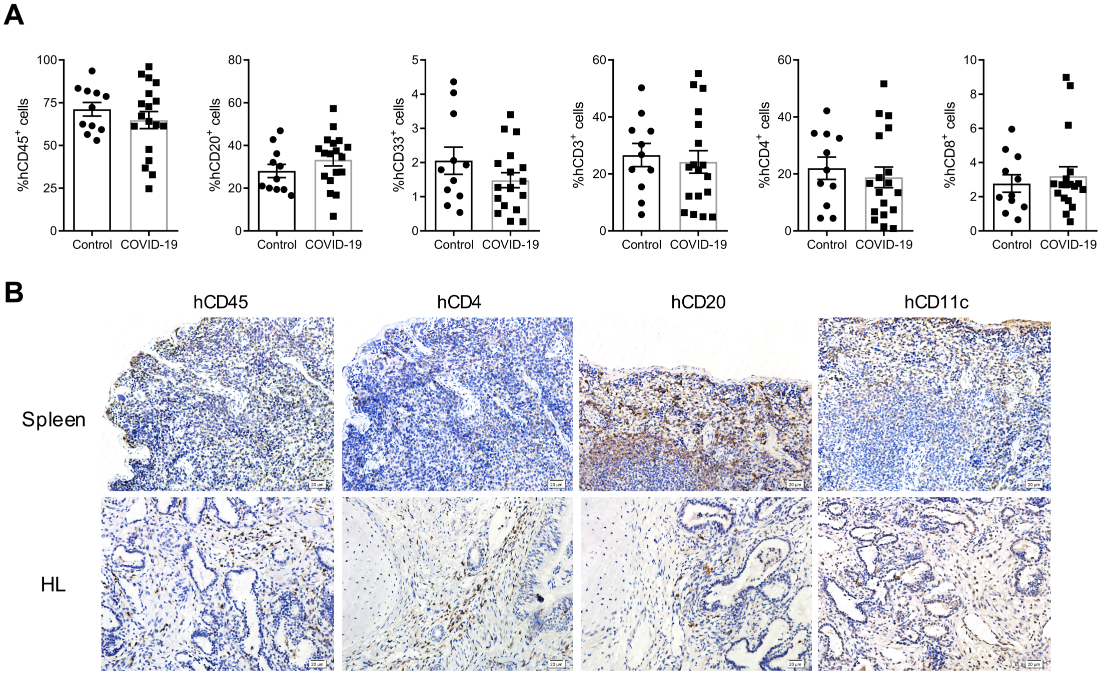


**Supporting Information Figure 2. Human immune reconstruction in HISL mice.** HISL mice were made by co-transplantation of human fetal lung tissue (subcutaneous) together with autologous human fetal thymic tissue (under renal capsule) and $CD34^+$ FLCs in NSG mice and used for examination. **(A)** Human immune cell chimerism in PBMCs of HISL mice at week 14. **(B)** IHC images of spleen (up) and HL (down) sections of HISL mice stained for human CD45, CD4, CD20, and CD11c.

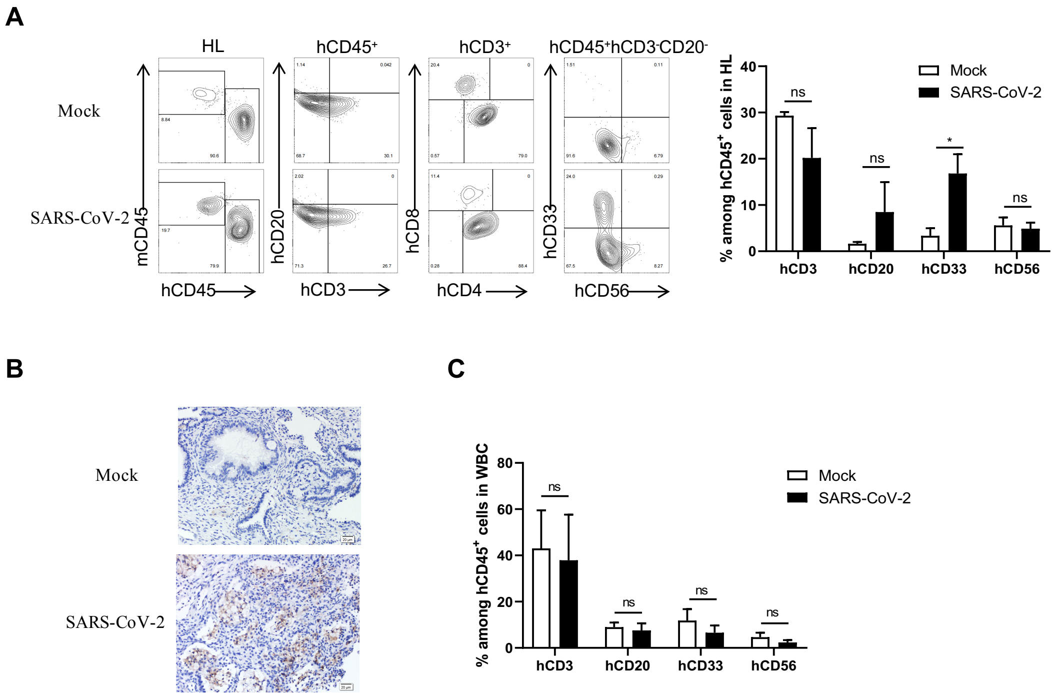


**Supporting Information Figure 3. Human immune cell profiles in HISL mice after SARS-CoV-2 infection.** HL graft and blood cells were prepared from SARS-CoV-2-infected (n=3) or mock-infected (n=3) HISL mice at day 21 and analyzed for human immune cells by FACS and IHC. **(A)** Representative flow cytometric profiles (left) and percentages (mean ± SEMs) of the indicated human immune cell populations (right) in HL cells. (**B**) IHC images of HL graft sections stained for human CD11c. (**C**) Percentages (mean ± SEMs) of the indicated human immune cell populations in WBCs from SARS-CoV-2- and mock-infected mice.

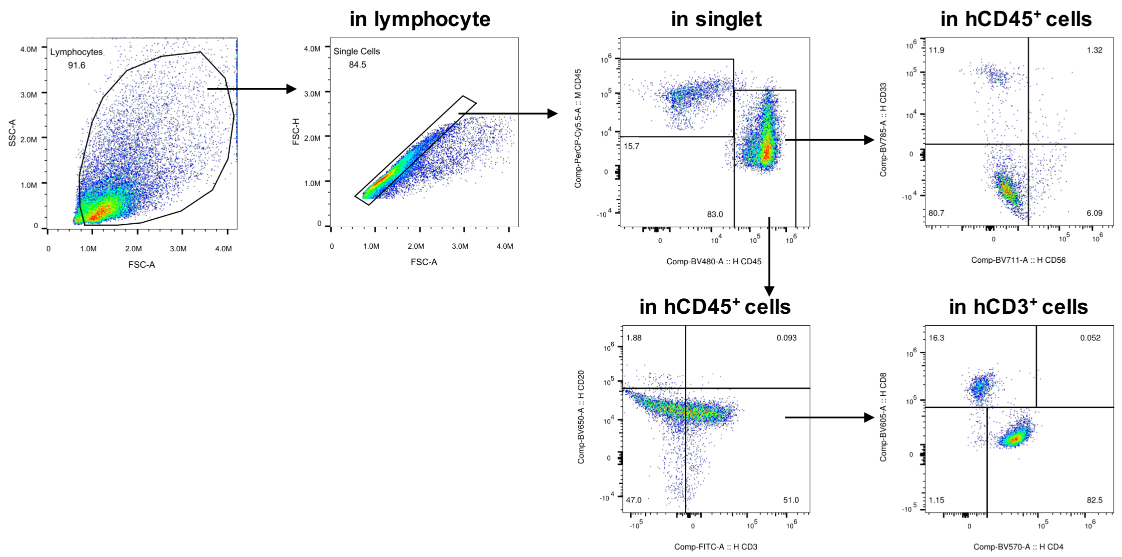


**Supporting Information Figure 4.** Flow cytometry gating strategies for experiments in Supporting Information Figure 2 and Supporting Information Figure 3.
